# Dynamics of individual visual biases in space and time

**DOI:** 10.64898/2026.06.22.733724

**Authors:** Mark Wexler

## Abstract

Recent work has brought to light a number of stimulus families whose perception is shaped by strong idiosyncratic biases. These biases differ significantly from one observer to the next, yet remain quite stable within observers when measured over multiple points in time, sometimes over months or even years. Nevertheless, we have previously shown that at least some of these biases undergo small but systematic changes over time. Although these temporal changes are also idiosyncratic, they generally act as a kind of memory that accumulates small random steps. Other research has shown that when stimuli are shown at different points in the visual field, biases can vary idiosyncratically across the spatial field as well. Here we ask whether variations in biases across space follow any regular pattern, and whether spatial and temporal variations are independent of one another. Measuring biases for surface orientation in structure-from-motion stimuli, sampled at numerous points in space and time, we find that variations both in time and space are positively autocorrelated: the closer two points are to each other in space or in time, the more similar the biases at those two points. We also found that spatial and temporal variations of bias are correlated, both between and within participants. Bias variations over space, time, and space-time are therefore not random but follow dynamics that may provide clues about the underlying mechanisms.

## Introduction

Individual differences in perception and cognition, in addition to their intrinsic interest, have traditionally been used as a tool to study the unicity or multiplicity of underlying mechanisms. If two effects have sizes that vary in the population, correlated variations argue for commonality of at least some of the underlying mechanisms (Spearman, 1904; Goodhew and Edwards, 2019; Brysbaert, 2024; Hedge et al., 2018). Recently, a different kind of individual difference in perception has come to the fore: large idiosyncratic biases in perception and action, with robust differences between observers (Deutsch, 1986; Carter and Cavanagh, 2007; Afraz et al., 2010; Mamassian and Wallace, 2010; Schwarzkopf et al., 2011; Houlsby et al., 2013; Schwarzkopf and Rees, 2013; Schütz, 2014; Wexler et al., 2015; Goutcher, 2016; Moutsiana et al., 2016; Schütz and Mamassian, 2016; Grabot and van Wassenhove, 2017; Kosovicheva and Whitney, 2017; Grzeczkowski et al., 2017; Witzel et al., 2017; Pressnitzer et al., 2018; Visconti di Oleggio Castello et al., 2018; Wexler, 2018; Lebovich et al., 2019; Drissi-Daoudi et al., 2020; Finlayson et al., 2020; Grabot and Kayser, 2020; Hwang and Schütz, 2020; Wang et al., 2020; Boeykens et al., 2021; Cretenoud et al., 2021; Van Geert et al., 2022; Wang et al., 2022; Wexler et al., 2022; Emery et al., 2023; Lebovich et al., 2025a). When measured over several points in time, these biases undergo changes that are typically smaller than differences between observers (Cretenoud et al., 2021; Drissi-Daoudi et al., 2020; Finlayson et al., 2020; Kosovicheva and Whitney, 2017; Wang et al., 2020; Wexler, 2018; Wexler et al., 2015). Some biases also show variations over space, that is over the visual field, and the spatial patterns of bias also seem to be idiosyncratic: they vary considerably from one observer to the next, but seem to be largely conserved over multiple measurements within observers (Afraz et al., 2010; Boeykens et al., 2021; Finlayson et al., 2020; Kosovicheva and Whitney, 2017; Moutsiana et al., 2016; Visconti di Oleggio Castello et al., 2018; Wang et al., 2020).

However, little attention has been paid to the systematic details of the spatial variations of the bias patterns. For example, if we know an individual’s bias at one point in the visual field, what if anything can we predict about nearby points in space? The same can be said about variations over time. The only studies to look at temporal variations in a detailed way have found that some biases act as a kind of stateful memory, accumulating changes over time (Wexler, 2018; Wexler et al., 2015). Finally, nothing at all is known about the spatio-temporal dynamics of biases: does knowledge of spatial variability inform us about temporal variability and vice versa?

Not all individual variations in perception are the same. Some are variations in the bias towards ambiguous stimuli with, and of those, some show only small degrees of interindividual variation (Mamassian and Goutcher, 2001), whereas others show much larger variations (Wexler et al., 2015). Other biases concern different degrees of susceptibility to illusions (Cretenoud et al., 2021; Grzeczkowski et al., 2017), and seemingly unambiguous perceptual judgments (Kosovicheva and Whitney, 2017; Lebovich et al., 2025b). There have been demonstrations that some idiosyncratic visual biases are due to, or at least correlated with, some physiological features. For example, a study has shown that the structure of a human observer’s corpus callosum can predict bias in the perception of the ambiguous motion-quartet stimulus (Genç et al., 2011), and other studies have shown similar correlations between other brain structures and other idiosyncratic biases in vision (Moutsiana et al., 2016). Surely then these physiologically-based biases cannot vary dynamically over time, at least not over short time scales? Surprisingly, in at least one case, the opposite has been found. I have previously shown that biases in the motion quartet, far from being fixed by the anatomical structure of the corpus callosum, can vary over time scales as brief as several hours (Wexler, 2018). Perhaps the bias in this case is determined by a combination of anatomical features and internal dynamic variables that can fluctuate. This possibility opens the door that many biases, even ones thought to be tied to anatomical features of the brain, may admit fluctuations.

This study made use of ambiguous structure-from-motion (SFM) stimuli, whose perception has previously been shown to depend on strongly idiosyncratic biases for surface orientation. These biases are measured from brief presentations of ambiguous optic flow that is compatible with two opposite surface tilts that vary from trial to trial (see Figs. 1a-c). These biases in the initial perception of ambiguous stimuli are therefore analogous to so-called onset biases in binocular rivalry (Carter and Cavanagh, 2007), and if the stimulus were to be presented longer it would undergo perceptual oscillations typical of bistable stimuli (Brascamp et al., 2018). These biases vary considerably from one observer to the next, that are quite stable over time, but when measured carefully the time series are found to undergo systematic accumulation of changes (Wexler et al., 2015). In this study I measured these biases in enough spatial locations and points in time in order to be able to draw novel, quantitative conclusions about the detailed spatial and spatio-temporal dynamics of perceptual biases.

**Figure 1.**
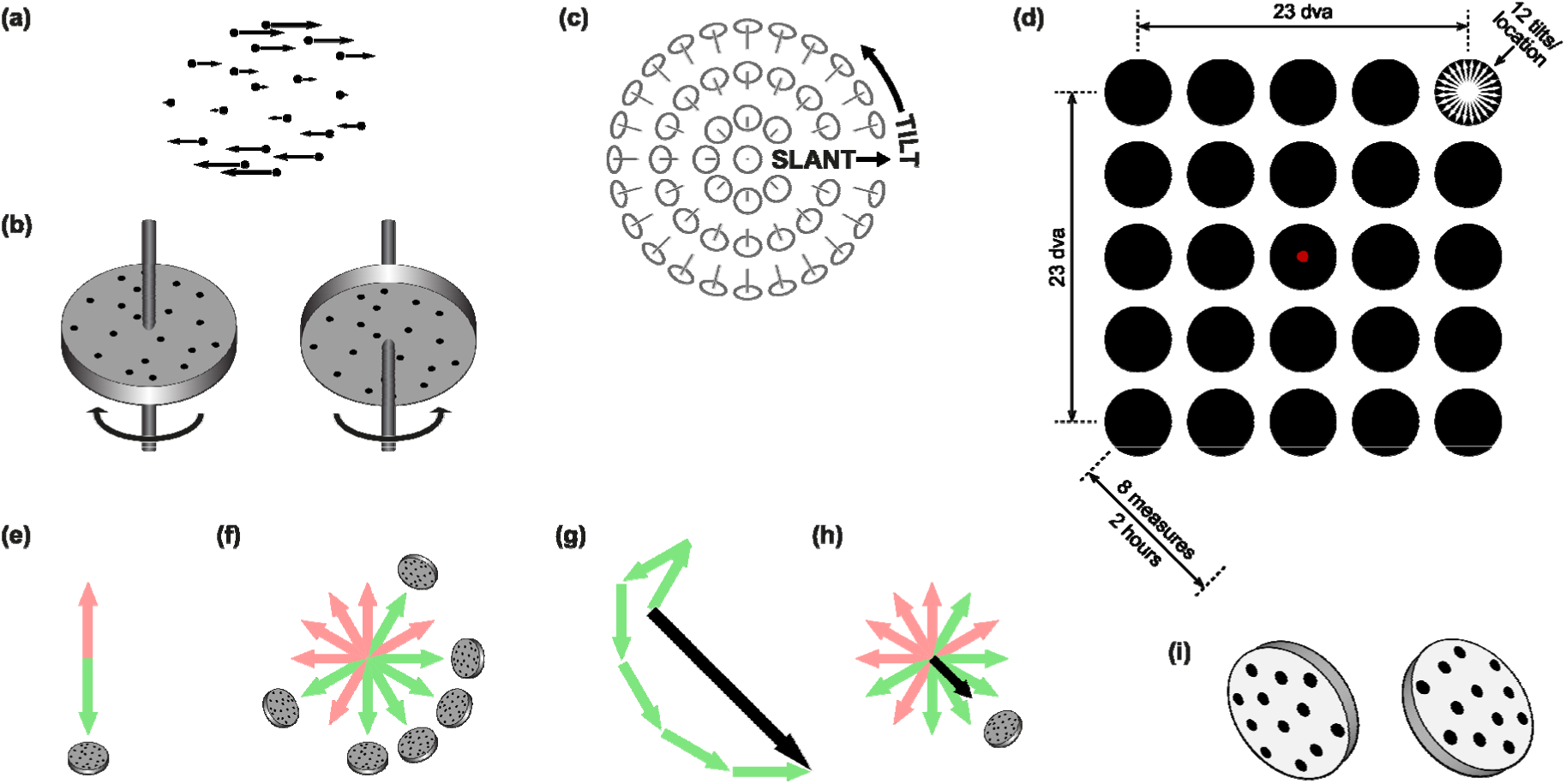
Stimuli, conditions, and basic data analysis. (a) Stimuli are sets of dots undergoing optic brief motion on a computer monitor. In this example, the arrows show the direction and speed of each (b) This stimulus is usually perceived as a surface slanted in depth undergoing a small rotation, but because of the properties of first-order optic flow, the orientation and direction of motion of the surface are ambiguous. In this example, two opposite orientations can be perceived (each with a different motion direction). The participants’ task was to indicate which of the two orientations they perceived. (c) 3D surface orientations can be parametrized by two angles, called slant and tilt. In our case, the tilt is ambiguous. All stimuli correspond to one of two tilts that differ by 180°. (d) A schematic of the experimental design. Stimuli were shown at one of 25 location on a virtual 5x5 grid, while participants fixated in the center of the monitor. At each location the stimuli were oriented at one of 12 tilts (each shown as 2 opposite-facing arrows). The locations and tilts were presented in random order. Each such block of 300 trials lasted on the average 12-13 minutes. Eight blocks were carried out, with each block starting every 15 minutes. (e) On each trial, one of two opposite tilts was reported. In presenting these responses, the trial will be shown as two opposite arrows corresponding to the two opposite tilts. The reported tilt direction is shown in green, the opposite, unreported tilt in pink. (f) An example of 6 trials in which the tilts sample 360°. The reported surface orientations are shown in higher contrast around the rim of the circle, and the corresponding tilts as green arrows. (g) In order to determine the bias, a vector mean is calculated from the reported tilts. (h) The resulting bias is shown as a black arrow. Its direction gives the preferred tilt, and its length (normalized to run from 0 to 1) measures the strength of the bias. (i) An example of a probe, here following a trial with tilt 37.5°, equally compatible with tilt 217.5°. The participant chooses which of the two tilts is closest to the perceived tilt. The left or right placement of the two possible tilts on each trial was randomized.

## Methods

Twenty-two participants viewed brief ambiguous structure-from-motion (SFM) stimuli at 25 locations on a virtual 5 x 5 grid, spaced by 5.7° of visual angle vertically and horizontally (Figs. 1a-d). Each stimulus was perceived as a rotating surface inclined in depth. Stimuli at each location were shown at one of 12 orientations; because of the parallel projection that was used, each surface orientation was compatible with one of two tilts that differed by 180°: 7.5° or 187.5°, 22.5 or 202.5°, …, 172.5° or 352.5°. Thus, 12 different stimuli were shown at 25 different spatial positions, for a total of 300 conditions per block. The trials were shown in random order, both with respect to position and orientation. The participant’s task was to fixate the center of the display (gaze monitored by an eye tracker), and following the presentation of each stimulus to report which of the two possible tilts was perceived. A single block took on the average around 11 minutes. Eight blocks were performed in total, starting approximately every 15 minutes. From the participant’s responses on the 12 trials at each position in each block, a bias was calculated (Figs. 1e-h). Thus a field of biases was measured at each of 5 x 5 spatial positions every 15 minutes, for a total of 8 measurements over time, which I will call time slices.

### Stimuli

Stimuli were generated by first placing dots on uniformly chosen random positions on a disk of radius 3.5 cm, with a density of 25 dots/cm^2^. (All spatial dimensions will be given in cm on the plane of the monitor, approximately 180/*π* ≈ 57.3 cm from the participant’s right eye, thus 1 cm is approximately 1 degree of visual angle.) The dot positions were then rotated about the origin in order to lie on a plane with slant *σ* = 45° and tilt *τ* = 7.5°, 22.5°, …, or 172.5° (12 equally spaced values, equivalent to 187.5°, 202.5°, …, 352.5° because of the symmetry in the orthographic projection: see below). We define slant *σ* and tilt *τ* of a plane so that the plane’s normal is equal to the vector (sin *σ* sin *τ*, sin *σ* cos *τ*, cos *σ*), where in our reference frame the horizontal *x*-axis points to the participant’s right, the vertical *y-*axis points upwards, and the *z-*axis towards the participant. The origin of the reference frame corresponds to the center of the monitor. Angles are defined so that 0° is towards the positive *x*-axis (to the participant’s right), and positive angles are counterclockwise from the participant’s point of view. In order to place the dots on a plane with slant *σ* and tilt *τ*, they were first rotated about the positive *y-*axis by angle *σ*, then about the positive *z*-axis by angle *τ*. This defines the base orientation for each stimulus.

The dots were always shown in motion, for a duration of 0.5 s. The motion corresponded to a rotation of the dots positions about the axis *â* = (cos(*τ* + 45°), sin(*τ* + 45°), 0), at a speed of 30°/s, centered on the central orientation (see above). Thus, the orientations shown corresponded to rotations from -7.5° to +7.5° about axis *â*, of the base orientation of each stimulus.

Each frame was placed so that the center of the stimulus (*x*_0_, *y*_0_) fell at one of 25 locations, on a 5 x 5 grid in the plane of the monitor. The grid was spaced by 5.75 cm in both horizontal and vertical dimensions. Thus the *x*_0_ and *y*_0_ coordinates of the stimulus center locations were -11.5, -5.75, 0, +5.75, or +11.5 cm. The stimulus was placed in one of two ways, very similar but not identical. In one group of participants (Group 1), all dots were simply translated parallel to the monitor by vector (*x*, *y*, 0), where *x*, *y* were the coordinates of the stimulus center. In a second group (Group 2), the dots were rotated about position (0, 0, 180/*π* ≈ 57.3 cm) close to the participant’s right eye, so that the stimulus center fell on position (*x*, *y*, 0). Because *x*_0_ and *y*_0_ were considerably smaller than the distance from the eye to the monitor, these two placement operations produced nearly identical results.

Finally, each dot position was orthographically projected. In Group 1, the projection was parallel to the *z* axis: a dot at position (*x*, *y*, *z*) was projected to position (*x*, *y*) on the monitor. In Group 2, the projection was parallel to the line of sight, the line from the eye position (0, 0, 180/*π*) to the stimulus center (*x*_0_, *y*_0_, 0). Once again, because displacements of the stimulus centers *x*_0_ and *y*_0_ were considerably smaller than the distance from the eye to the monitor, these two projections produced nearly identical results.

In particular, all stimuli were ambiguous, compatible with either tilt *τ* or tilt *τ* + 180°. This is a property of so-called first-order optic flow, due to the orthographic projection that was used (Hoffman, 1982; Koenderink, 1986). In more detail: all stimuli were identical under the simultaneous transformation of the tilt *τ* ↦ *τ* + 180° and angular velocity ***ω*** ↦ -***ω***.

The dots were drawn as white disks of diameter 0.08 cm on a black background. A fixation point was drawn at the center of the monitor, i.e., position (0, 0), a red disk of diameter 0.2 cm.

Each trial was preceded by the fixation point displayed alone in the center of the monitor for 750 ms. During this phase the fixation flickered at 5 Hz. In order to prevent drowsiness, a brief auditory chirp was played on 20% of the trials chosen at random. This was followed by the 500 ms stimulus sequence, described above. During the stimulus the fixation dot remained in the center of the monitor, and did not flicker.

If the participant was deemed to have fixated the central fixation point (see below), the trial was concluded by a response phase, during which the participant indicated the perceived average tilt of the surface—average because the tilt actually varied by a few degrees in the course of the motion. The perceived tilt was reported by selecting one of two images displayed to the left and right of the center of the monitor. The images were unambiguous static depictions of surfaces with the two possible tilts on the given trial, namely *τ* or *τ* + 180°. An example for tilt *τ* = 37.5° is shown in Fig. 1i. The left or right placement of the two possible tilts on each trial was randomized. The participant selected the perceived average tilt by moving a cursor to the corresponding image and clicking a mouse button.

### Procedure

The experiment was performed in a nearly dark room. Participants were seated facing a 20 inch CRT monitor, with their chins on chinrest and foreheads against a forehead rest. The participant’s right eye was positioned approximately 57.3 cm from the monitor, horizontally and vertically centered on the monitor. The left eye was covered by an opaque patch.

The experiment began with a short training block in which the task was introduced to the participant. During the training the stimulus always appeared at the center of the monitor. The experimenter emphasized to the participant that during the main part of the experiment the stimulus would usually appear away from the central fixation point, which it was crucial to fixate.

To ensure fixation on the fixation point at the center of the monitor, participants’ right eyes were tracked using an EyeLink 1000 infrared video-based eye tracker. Following the stimulus of each trial, several checks were performed on the eye movement recording during the 500-millisecond duration of the stimulus:

- The mean gaze position had to less than 2.875 cm (2.87 degrees of visual angle) from the fixation point (half the distance between stimulus locations)
- There could be no saccades with amplitudes over 1 cm (1.00 dva)
- There could be no blinks

If any of these tests failed, the trial was scheduled to be repeated at a randomly chosen later point during the same block. If the eye tracker returned no samples during the stimulus, a new set-up and calibration was performed.

The main experiment consisted in 8 blocks of 300 trials each (referred to as “time slices” in the article). Each block was a factorial design, with 5 x 5 spatial locations and 12 tilts at each location. The order of trials was randomized both respect to location and tilt.

The duration of the 300 trials in each block varied. The distribution of the durations is shown in the left graph below. Median duration was 11 minutes and 15 seconds, with only 3 blocks out of 144 having durations above 15 minutes. As soon as a block was finished, the participant was told to rest for a duration calculated so that the next block would begin 15 minutes after the previous block, but always at least 30 seconds. When the rest period was over, participants began the next block, which always started with an eyetracker calibration. In order to measure the real time difference between blocks, I calculated the block asynchrony, defined as the time difference between the start of the first trial of a block and start of the first trial of the previous block. The actual distributions of block durations and asynchronies are shown in Fig. 2. While the goal was to have block asynchronies of 15 minutes, because of the time differences required in setting up and calibrating the eye tracker, blocks lasting longer than average, and participants occasionally requiring extra time between blocks, the exact asynchrony of 15 minutes could not be achieved. The median asynchrony was 15 minutes and 30 seconds.

**Figure 2.**
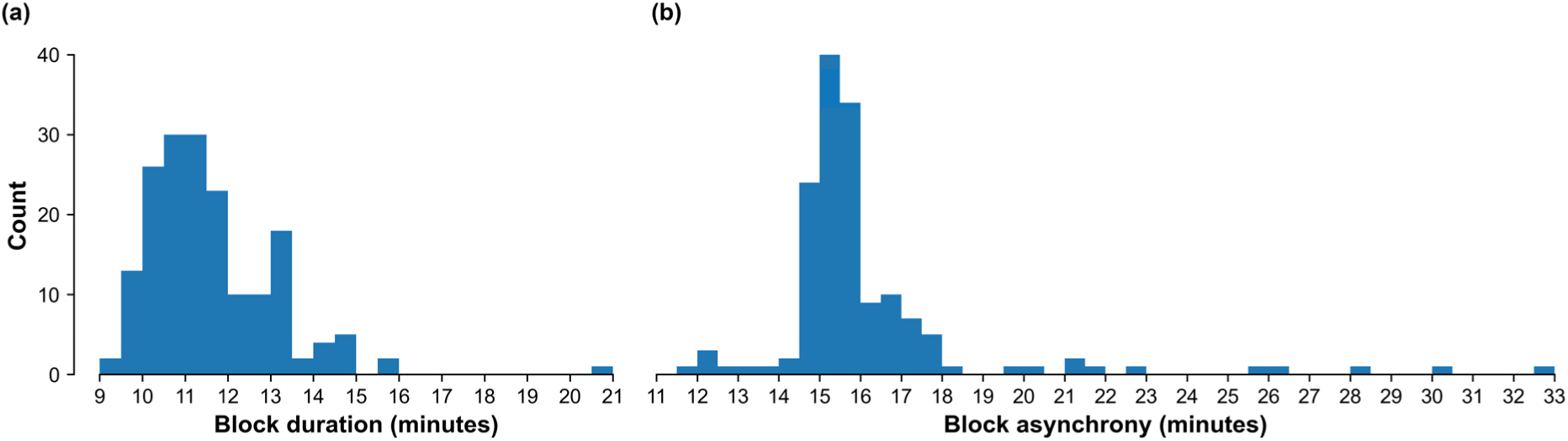
The timing of the experimental blocks. (a) The distribution of block durations. (b) The distribution of block asynchronies, i.e., the time differences between the start of successive blocks. The goal was to have block asynchrony of 15 minutes.

26 people with normal or corrected-to-normal vision participated in the experiment in exchange for payment of 10 €/hour. Participants were eliminated from further analysis for one of two reasons: either they had more than 2% of their eyetracker samples that went further from the central fixation zone, defined as the square centered on the central fixation point and whose side is equal to the spacing of the stimuli, or more than 20% of their trials were missing because of head motion during a session. Four participants were eliminated for these two reasons. In the remaining participants, 0.3% of the trials had gaze that went outside the central fixation zone (call these ‘erroneous trials’). The responses for these trials were replaced by one of the two tilts compatible with the corresponding stimulus that had a positive projection on the vector average of the remaining non-erroneous trials at that location and in that session (time slice). In total, 0.05% of the trials had their response flipped following this procedure.

There were 2 groups of participants: 14 in group 1 and 8 in group 2. The distinction between the groups was a small difference in the way the stimuli were projected (in group 1 the projection was orthogonal to the monitor, while in group 2 the projection was orthogonal to the plane perpendicular to the line between the eye and center of each stimulus). Because no significant group effects, were found, the data was collapsed across groups.

## Results

The bias is a 2D vector that measures the observer’s propensity to perceive tilt in a particular direction. It is defined as the mean of the unit vectors corresponding to each reported tilt (the unit vectors are shown as green arrows in Figs. 1e-h, the bias as the black arrow in Fig. 1h). When there is no bias, the perceived tilts are uniformly distributed around the 360° circle, and the resulting bias vector is short. On the opposite extreme, all perceived tilts lie in a particular 180° fan (±90° about a central tilt), and the bias vector has maximum length (normalized so that the maximum length is 1) and points towards the center of the fan. Thus the length of the bias vector measures the strength of the bias, and its direction measures the preferred tilt (see Wexler, Duyck & Mamassian (2015) for further details).

Figure 3 shows illustrative examples of partial times series for four participants, with raw responses for all spatial positions and the corresponding bias vectors. The raw data from all the 52,800 trials is shown in Fig. S1 in the Supplementary matrials. Several aspects of these data stand out. Biases are strong at most spatial locations at most times, but they do vary across space to different degrees, across time and between participants. Not only do biases differ at any given spatial location between participants, but overall spatial *patterns* of bias variation also show individual variation. The bias at a given location does not seem to be independent of biases at nearby spatial locations: on the whole, nearby biases seem to be more similar than biases father apart, although there are some exceptions. Finally, the spatial bias patterns remain similar over time, but with variations for different locations and different observers.

**Figure 3.**
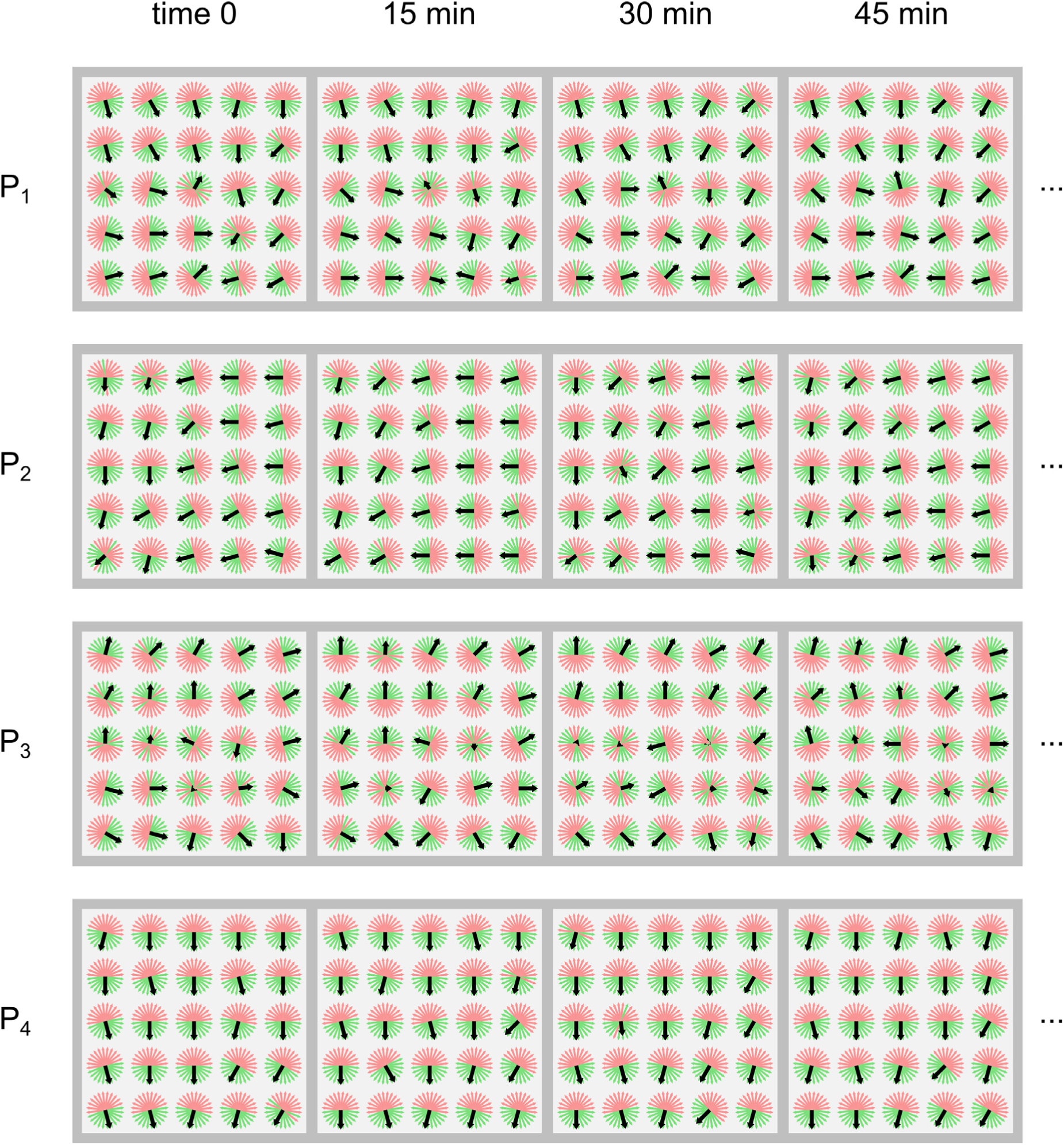
Examples of partial raw data for four participants. Each line shows four blocks (half the time series) for one participant. Blocks began every 15 minutes, so the data corresponds to about one hour. Each 5x5 square corresponds to one block or time slice. The 25 circles represent the data from the different spatial locations. The response for each of the 12 trials at each location is shown as two opposite arrows, with the green arrow corresponding to reported tilt, as described in Figs. 1e,f, and the pink arrow the opposite, unreported tilt. The black arrows show the resulting biases, calculated as shown in Figs. 1g, h.

### Are biases significant?

Do we have evidence of systematic biases? The *absence* of systematic biases would mean independent, random responses on individual trials, which would yield short bias vectors. I calculated the mean bias strength over the 25 locations at each time slice for every participant. This mean bias strength was compared to a null-hypothesis distribution, obtained by simulating unbiased observers (who select one of the two responses randomly with equal probability, independently on each trial), and calculating an empirical distribution from 10^5^ bias strengths, each averaged over 25 realizations of 12 trials. Mean bias strength was significant on every time slice in 20 out 22 participants, and in the remaining two participants in was significant in 4 and 6 times slices out of 8, after a Benjamini-Hochberg correction for multiple tests using a false discovery rate of 5%. Thus, there was a significant bias at least in some parts of the visual field at almost all times for all participants.

### Are spatial bias patterns idiosyncratic?

A simple definition of idiosyncraticity of a repeatedly measured quantity is whether within-participant variations of the repeated measurents are lower than variations between measurements in different participants (used for example by Kondo, Murai & Whitney, 2022). I defined the difference between two spatial patterns of biases as the mean Euclidean distance between the biases at each of the 25 positions. For each participant I then calculated the mean difference between all pairs of his or her own time slices of spatial bias patterns (‘within’), and the mean difference between other participants’ time slices (‘between’). The results can be seen in Fig. 4, which also shows the standard deviations of these variability measures. The mean within-participant difference was 0.36, and the mean between-participant difference was 0.78. Moreover, the between-participant difference was higher than the within-participant difference for all 22 out of 22 participants. A paired t test showed that the between-participant differences were significantly higher than the within-participant ones (*t*_21_ = 14.7, *p* < 0.0001). We can therefore conclude that time slices of spatial bias patterns resemble each other more within individuals than across individuals—in other words, the spatial bias patterns are idiosyncratic.

**Figure 4.**
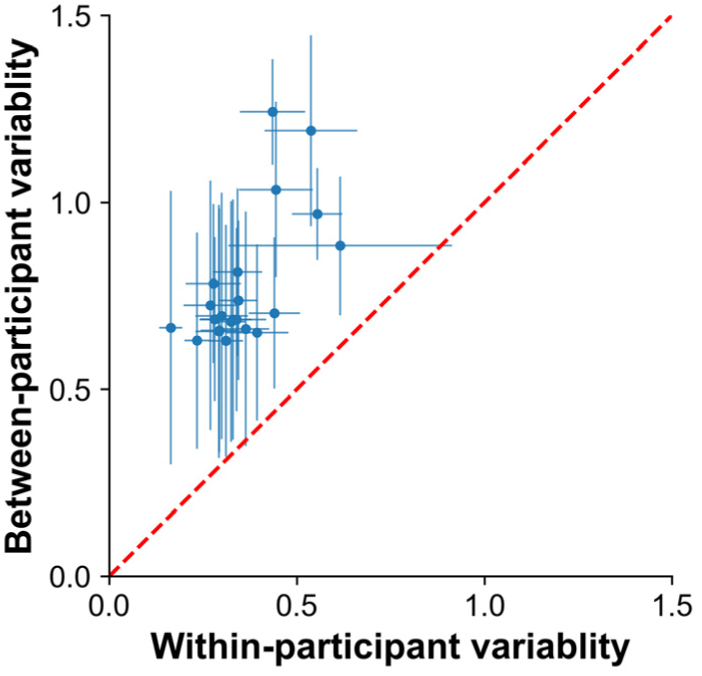
Between- versus within-participant variability of spatial bias patterns, demonstrating their idiosyncraticity. Variability is defined as the mean Euclidean distance between bias vectors at corresponding spatial locations, averaged over all pairs of time slices either within one participant, or between a participant and all other participants. Each dot corresponds to one particpant, with the error bars showing the standard deviations over pairs of time slices.

An alternative way of addressing the same question is to ask if there is something sufficiently idiosyncratic about individuals’ spatial bias patterns so that a machine-learning algorithm can learn to identify participants from their spatial bias patterns. In particular, I taught a support-vector classifier with a third-order polynomial kernel (SVC from the scikit-learn package) to predict participants from time slices of their spatial bias patterns. (Several other classifiers gave similar results.) The classifier was trained on 7 out of 8 time slices, and tested for generalization on the remaining slice. (The two cartesian components of the bias vector were given as separate predictors at each spatial position, yielding 2 x 5 x 5 = 50 separate predictors.) Generalization performance—the percentage of participants that were correctly predicted—was 94.3% (averaged over the 8 time slices). Thus individual spatial bias patterns permit very precise identification of the individual, even on unseen patterns, and are therefore idiosyncratic.^1^

### Spatial properties of biases

#### Do biases vary across space?

As can be seen in Figs. 3 and S1, biases often varied across spatial locations in the visual field, but did so to different extents in different participants and at different times. To check whether some of the spatial variation was significant, I calculated the largest difference between any pair of 25 biases in any single time slice, for every participant (expressed as the square of the Euclidean distance between vectors). These maximum differences were compared to null-hypothesis distributions of the same measure, based on the assumption that the biases at all 25 spatial locations were derived from the same distribution. This null distribution was derived empirically for each time slice in each participant, by averaging the 25 sets of responses after rotating them so that bias direction at each location was in the same direction. The probabilities of the null hypothesis were calculated by converting the actual maximum bias differences to quantiles of the null-hypothesis distribution derived from 10^5^ samples calculated using the above procedure, and correcting for multiple tests using the Benjamini-Hochberg procedure with false discovery rate 5%. I found that there was significant variation of bias across space in at least one time slice in all 22 out of 22 participants, with significant variation at *all* time slices in 16 out of the 22. Therefore biases do vary across space, in most participants most of the time.

#### Are nearby biases correlated?

Studying the examples shown in Fig. 3, it becomes clear that most bias vectors are close to the biases at neighboring positions, at a given time. This property is known as *spatial autocorrelation*: values of a field are correlated with values at nearby points (Cressie, 1993). Spatial autocorrelation can be thought of as a discrete stochastic generalization of the notion of continuity for a field.^2^ One way of visualizing spatial autocorrelation is through Moran’s diagram (Bouayad-Agha and Bellefon, 2018), in which the values of a field at each sampled point are shown on one axis, and the mean values of its nearest neighbors are shown on the other axis.^3^ Such a graph, for the eight time slices of a representative participant, is shown in Fig. 5a. The correlation that is evident in Fig. 5a can be quantified by an index known as Moran’s *I* (Bouayad-Agha and Bellefon, 2018), close to Spearman’s correlation for the points shown in Fig. 5a. I calculated this index for each time slice for every participant, separately for the horizontal and vertical components of the bias, and averaged the Fisher z-transformed indices weighted by the variance of each time slice.^4^ The values of Moran’s I for individual participants ranged from to −0.01 to 0.74 for the horizontal component of the bias, and from −0.06 to 0.70 for the vertical component. The significance of these spatial autocorrelations was evaluated by using a randomization test, in which the biases were scrambled in space and time. This test (with 10^4^ randomizations per participant, and Benjamini-Hochberg correction with false discovery rate 0.05) revealed that all 21 out of 22 participants had significantly positive spatial autocorrelations for the horizontal component of bias, and 18 out of 22 did so for the vertical component of bias. Combining individual autocorrelations, again using z-transformed Moran’s I indices weighted by participants’ overall variance, I found average autocorrelations of 0.62 and 0.46 for horizontal and vertical bias components, respectively. A randomization test (10^6^ randomizations) revealed that the grand average spatial autocorrelation was significantly positive for both horizontal and vertical components of bias (*p* < 10^-4^).

**Figure 5.**
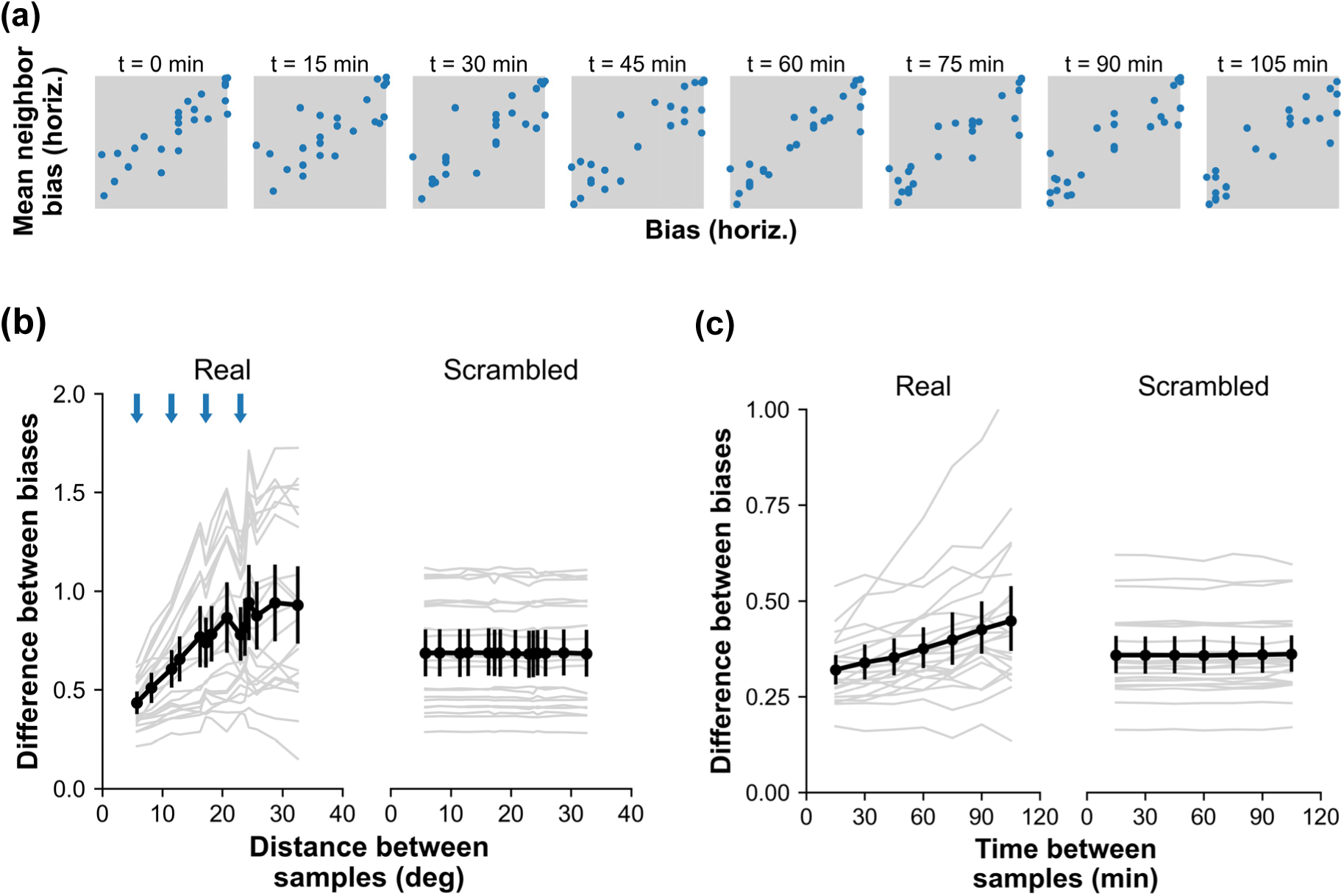
Spatial and temporal autocorrelation of biases. (a) Moran’s diagrams showing spatial autocorrelation for a representative participant, for the horizontal component of the bias vector. At each of the 8 time slices, every point represents the bias on the *x* axis, and the mean of its nearest spatial neighbors on the *y* axis. The strong correlation shows that there is significant spatial autocorrelation, meaning that nearby locations have closer biases than locations farther apart. (b) In the “Real” graph on the left, curves that show the differences in pairs of biases sampled at all pairs of spatial locations (not just nearest neighbors) but in the same time slice. The horizontal axis shows the spatial distance between the two samples, while the vertical axis shows the mean Euclidean distance between bias vectors sampled at that distance. The gray curves represent individual participants (averaged over time), while the black curve shows the mean over participants (the bars represent the 95% confidence interval of the mean). The 4 blue arrows show distances corresponding to 1, 2, 3, and 4 grid spacings. The “Scambled” graph on the right shows the same analysis but with the 25 spatial positions scrambled (mean over 100 scramblings). (c) Same analysis as in (b) but over time, showing how difference between biases measured at the same spatial location but at different times depends on the time between measurements. The gray curves represent individual participants (averaged over time, and binned by distance) while the black curve shows the mean over participants and 95% confidence intervals. The “Real” graph shows the analysis for actual data, the “Scrambled” graph shows the same analysis but with the 8 time slices scrambled (mean over 100 scramblings).

Another way of visualizing spatial autocorrelation is through a graph that shows, for each pair of biases sampled as different positions in space (but in the same time slice), the mean difference between the biases, as a function of the distance between the two positions (similar to the variogram commonly used in spatial statistics: see Cressie, 1993). For fields that are spatially autocorrelated or continuous, this distance-difference curve should be smaller for smaller distances and larger for larger distances. (If the field is uniform over space, than the distance-difference curve will be flat.) The curves in the “Real” graph of Fig. 5b show mean Euclidean difference between bias vectors (since bias vectors fall inside the unit disk, the Euclidean difference runs between 0 and 2) as a function of distance between positions, for individual participants and a mean over all participants. Performing a linear regression of mean difference versus distance, I found that the regression slopes of all 22 participants were positive. A separate analysis revealed that participants whose distance-difference curves had shallow slopes were those who had less variation in bias over space, as expected.

Carefully examining the individual participants’ curves in Fig. 5b reveals two points where many participants’ curves simulatenously dip. These points correspond to distances equal to 3 and 4 times the grid spacing (the blue arrows at the top of the graph mark 1-4 times the grid spacing).

At these values, the pairs of locations at which we calculate bias differences have purely horizontal or vertical separations. Although generally the difference between biases grows with the distance between measurements, purely horizontal or vertical separations seem to give rise to smaller differences than diagonal separations. The mean difference at distance 23 (corresponding to 4 grid spacings, and therefore horizontal or vertical separations, at the rightmost blue arrow in Fig. 5b) is 0.78, while the mean difference at the next smaller distance, 20.73 (corresponding to diagonal separations), is greater rather than smaller: 0.87. The difference between the two is significant (2-sided t test, *t*_21_ = 3.02, *p* < 0.01). The other visible dip, at 3 grid spacings, does not attain significance. If this effect holds up, it may indicate that changes in bias are smaller across purely horizontal or vertical separations than across diagonals.

The “Scrambled” graph on the right shows that the spatial autocorrelation of biases, or the fact that biases are closer to one another at positions closer to each other in space, depends on the actual metric structure of space. In the analysis shown in the scrambled graph, the spatial labels of the 25 positions were scrambled, so that their ‘distances’ no longer reflected the normal Euclidean metric of space. The curves show means over 100 such scramblings. As expected, the resulting distance-difference curves are flat: biases sampled at points ‘closer’ or ‘father’ away were actually sampled at arbitrary points in space.

These results on Moran’s I and distance-difference curves show that biases have the spatial structure typical of continuous or spatially autocorrelated fields: the closer two points are in space, the more similar the biases at the two points.

#### Is there a central tendency in the spatial patterns across participants?

Although the focus of this study is individual biases, it is interesting to check whether individual patterns bias patterns have a central tendency. To answer this question, I calculated the mean bias vector at each location, averaging over participants and times. The resulting grand spatial mean pattern of biases, shown in Fig. 6a, is extremely suggestive, as it resembles the spatial pattern of surface norms of a ground plane seen from above.

**Figure 6.**
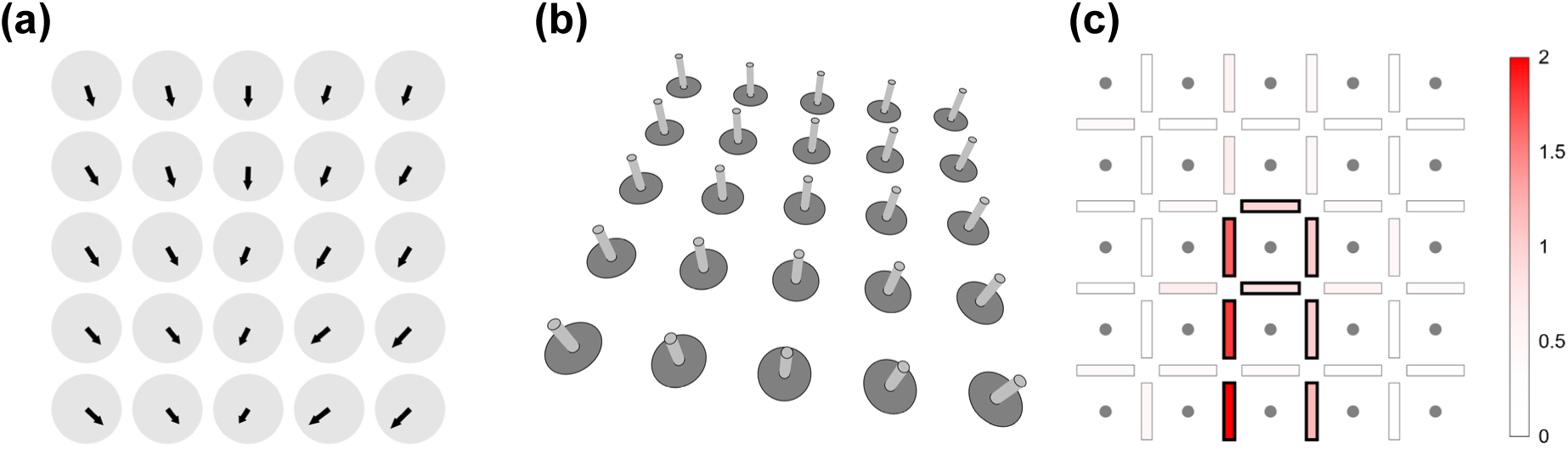
Global spatial patterns of bias. (a) Mean bias at each spatial position, averaged over all time slices and participants. While this bias is weaker than bias calculated for individual participants at particular time slices, it consitutes a significant central tendency as a spatial pattern. (b) When the mean biases are considered as normal directions of a projected planar surface, the plane closest to the mean bias pattern, pictured here, has slant 39.3° and tilt 82.3°, close to the ground plane seen from above. (c) A map that shows bias changes between nearest neighbors on the 5x5 grid of positions. The gray dots represent positions, and the rectangles between each pair of nearest neighbors represent the mean absolute bias difference, with greener shades corresponding to larger differences. The rectangles corresponding to the 20% of the pairs having the largest difference are outlined in black.

Our previous study (Wexler, Duyck & Mamassian, 2015) had already shown that, while in the center of the visual field biases occur at all tilt directions, on the population level the most common bias corresponds to the tilt of the ground plane. It is notable that the central tendency of the bias field shown in Fig. 6a seems to reproduce the effects of perspective in viewing a ground plane: the biases tilt to the left on the right side of the visual field and vice versa on the left side; and they tilt more at the bottom of the image (corresponding to the parts of the ground plane closer to the observer) than at the top. In fact, the effects of perspective are exaggerated: they would not as strong for a surface whose size was only 23° of visual angle. I fit a plane to the mean bias map, finding a plane whose normals were as close as possible in their directions to the mean biases, varying both the slant and tilt of this plane, as well as a scale factor to account for the exaggerated perspective. The result was a plane shown in Fig. 6b, having slant 38.5° (close to the simulated slant of 45°) and tilt 263.1°, close to the tilt of the ground plane (270°).

Because of the robust central tendenency present in the biases, an argument can be made that the relevant spatial degree of freedom in the biases isn’t the raw pattern of variation across space in each participant, but the difference between that pattern and the grand spatial mean map (Fig. 6a): the variations in the biases about the grand spatial mean. Removing the central tendency reduces the variance of the biases by a mean of 32.2%. It is notable that the analyses of spatial autocorrelation given above (the distance-different slopes and Moran’s I) still yield significant spatial autocorrelations when applied to the variations in the biases about the grand spatial mean.

#### Where in the visual field do biases vary the most across small distances?

If biases at neighboring positions tend to be close to one another, is this difference between neighbors uniform across the visual field? To answer this question, I calculated the Euclidean difference between biases at each pair of nearest-neighbor positions, and then averaged over times and participants. The result is shown in Fig. 6c, using shading to represent average variations across neighbors. The variations are far from uniform, and are largest across the vertical midline in the lower half of the visual field. (Because the the middle column falls exactly on the midline, variations across the midline may fall on either side of it.) The large changes in bias across the lower midline can also be seen from the mean bias map in Fig. 6a. Testing the statistical significance of this particular spatial pattern is complicated by the danger of ‘double dipping’ (Kriegeskorte et al., 2009), using the same data to both localize and test the zone of highest variation. In order to avoid double dipping, I used the leave-one-out technique: for each participant, the data of the remaining participants was used to find the 20% of the neighbors having the highest variation in bias (call them the “hot pairs”); then, for the left-out participant variation was calculated across those hot pairs, and across the other 80% (the “cold pairs”). Repeating this procedure for each participant, I found that the mean change in bias (using the Euclidean distance between bias vectors) across the hot pairs was 0.61, while across the cold pairs it was 0.39; a bootstrap test across participants revealed that this difference was significant (*N* = 10^6^, *p* < 10^-6^). The 20% of the most common hot pairs across participants are highlighted in Fig. 6c with black outlines. Thus the largest variation of biases over the visual field occurs at the center and around the lower vertical midline.

#### Do biases show a tendency to mirror symmetry?

The central bias pattern shown in Fig. 6b exhibits a mirror symmetry: biases at locations on the right of the visual field have a leftward horizontal component, while biases on the left have a rightward component. (Curiously, the biases on the lower midline tend to point leftwards.) This tendency is observed—less clearly of course—in individual bias patterns as well (Figs. 3 and S1), especially if we subtract the mean bias at each time slice. I performed a test for this mirror symmetry in individual data, by counting the proportion of the biases in the two leftmost columns that had a horizontal component in the opposite direction to the corresponding location on the right side of the visual field (individually for each participant and each time slice, after subtracting the mean bias). Averaging these mirror-symmetry proportions over all the time slices, I found that 21 out of 22 participants had a net mirror symmetry.

### Temporal properties of biases

#### Do biases vary over time?

As can be seen in Figs. 3 and S1, there were different degrees of temporal variation in different participants and different spatial locations. To check whether there were significant temporal variations in some cases, I applied the same maximum bias difference procedure as for variations over space (see “Do biases vary across space” above), but instead calculated the maximum bias difference over the 8 time slices at each spatial location for each participant. This test revealed that 20 out of 22 participants had significant time variations in bias in at least one spatial location (in 1 to 11 out of the 25 locations). It should be kept in mind that this is a minimum: there is probably more temporal variability that does not survive significance test. Thus biases do vary across time in at least some spatial locations in most participants.

#### In which directions do biases change over time?

In order to check if there were any preferred directions in bias changes over time, I calculated the bias vector steps over one time slice (i.e., 15 min) at the same spatial location. Fig. 7a shows the distribution of the directions of these steps (taking steps of length greater than zero), for all times, locations and participants. The smoothed circular histogram is fairly uniform, with weak peaks in the horizontal directions. A more informative way of looking at bias step directions is to take the initial and final bias in each pair to be subtracted, and to first rotate them so that the initial bias points to the right, that is along angle 0°. This gives a different meaning to the direction of the resulting bias step. If it is to the right (near 0°) then the bias vector becomes longer, or, in other words, the bias becomes stronger. If near to 180°, then, on the contrary the bias becomes weaker. If the step is near 90° or 270°, then the bias rotates counterclockwise or clockwise, respectively. The distribution of these rotated directions, shown in Fig. 7b, is highly non-uniform. There are two large peaks around 90° and 270°, corresponding to bias rotations, and two smaller peaks near 0° and 180° corresponding to increases and decreases in bias strength. This rough pattern is observed in nearly all individual participants. The rough mean proportions are around 70% of the steps as rotations in the clockwise or counterclockwise directions, with an approximate balance between the two directions; and 20% and 10% of the steps in the direction of weaker and stronger bias, respectively.

**Figure 7.**
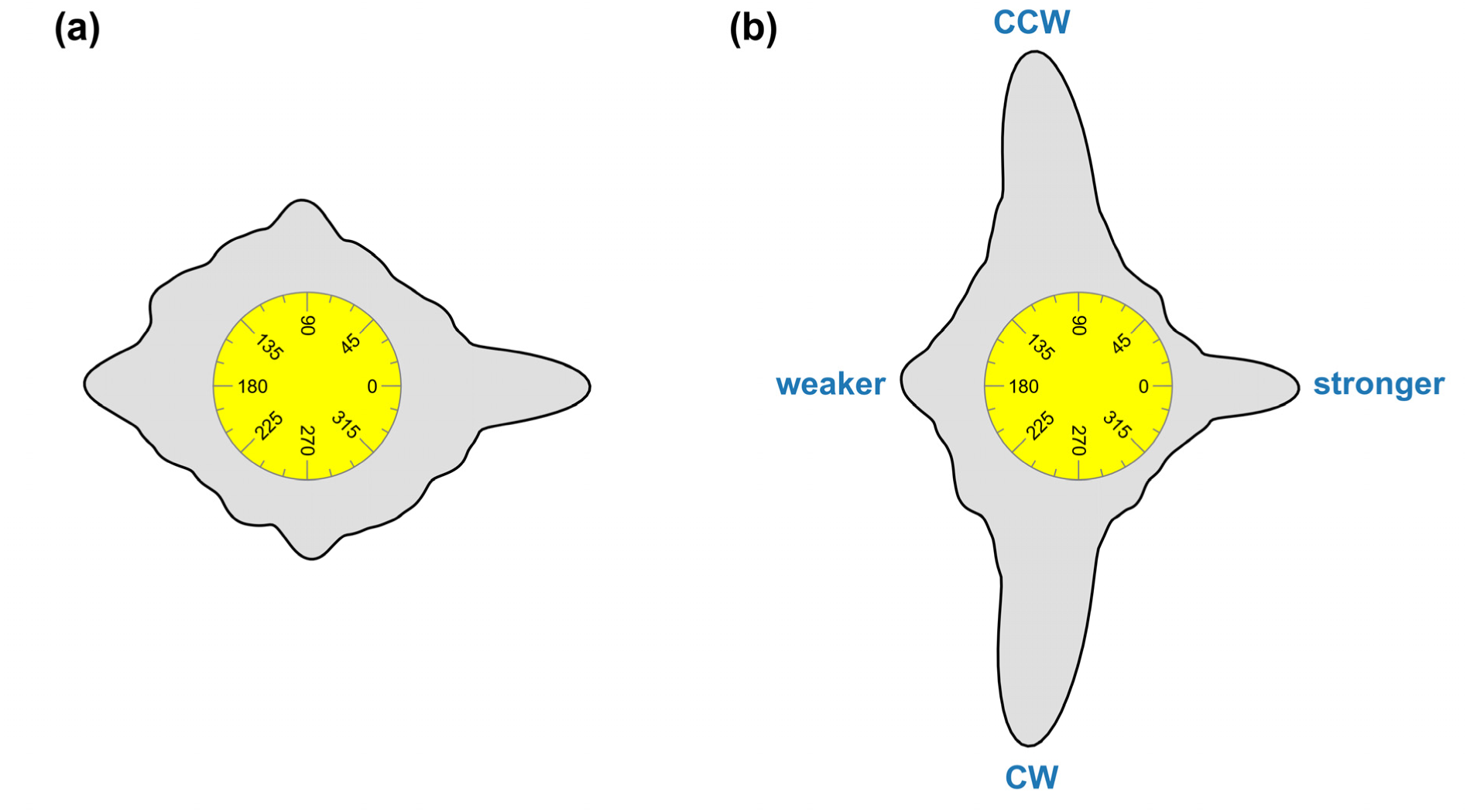
The directions of bias change over time. (a) A circular histogram showing the distribution of bias direction changes over one time slice (15 min) and at the same spatial location, over all spatial locations and participants. In each direction, the radial distance between the rim of the yellow circle, which represents zero, and the outer black curve represents the prevalence of bias changes in that direction. The individual directions were smoothed using the quartic biweight kernel with bandwidth 15°. (b) Similar analysis as in (a), but with all directions rotated so that the initial bias vector of each pair points to the right (i.e., has angle 0°). In this representation, angles around 0° and 180° represent increases and decreases in bias strength, respectively, while angles near 90° and 270° represent counterclockwise and clockwise rotations of the bias vector.

#### Are biases close in time correlated?

Similarly to the spatial version of this question discussed above, we can ask whether two biases, sampled at the same point in space but at two different times, are closer to each other when the two samples are closer in time. I first calculated an autocorrelation—here just the ordinary one-dimensional autocorrelation—of the bias time series of eight terms at each of the 25 spatial locations for each participant, and then took the mean of the 25 z-transformed autocorrelations weighted by their variances, to produce a single temporal autocorrelation estimate for each participant. These estimates ranged from −0.26 to 0.65 for the horizontal component for the bias, and from −0.31 to +0.41 for the vertical component. A randomization test like the one for the spatial autocorrelations (10^3^ randomizations per participant) revealed 9 out 22 participants had a significantly positive autocorrelation for the horizontal component of bias, and 7 out of 22 did so for the vertical component. I then calculated a ‘grand-average’ autocorrelation by averaging the participants’ autocorrelations and weighting by variance, and obtained +0.17 for the horizontal component and +0.05 for the vertical. Randomization tests on these grand averages (10^3^ randomizations per participant) revealed that both horizontal and vertical components of the bias had significantly positive temporal autocorrelations (*p* < 10^-3^).^5^

In order to analyze the temporal structure of the time series at lags above 1 time slice (15 minutes), in the “Real” part of Fig. 5c I plotted the average Euclidean difference between biases at the same spatial position as a function of time between samples. Regressing the individual-participant curves against time revealed that 20 out of 22 had positive slopes. Comparing to similar curves for the spatial autocorrelation (Fig. 5b) shows that biases accumulate changes over both space and time in most cases, but changes over 2 hours are smaller than changes over 23 degrees of visual angle. The “Scrambled” part of the figure shows that the time labels are randomized so that time no longer has the standard metric structure, the distance-difference curves become flat, as they do for space (Fig. 5b).

### Spatio-temporal connections

Now that we have examined the purely spatial of the biases across the visual field and their purely temporal properties across time, we turn to the interactions between the variations across space and across time.

#### Do participants who have larger spatial variations in biases also have larger temporal variations?

Consider Fig. 8a, which shows the variability of the biases across space and time in each participant. By careful inspection it can be seen that participants who have large variability across time (i.e., large differences in the transparent arrows at single locations) are by and large also the ones who have large variability over space. The following analysis formalizes this observation.

**Figure 8.**
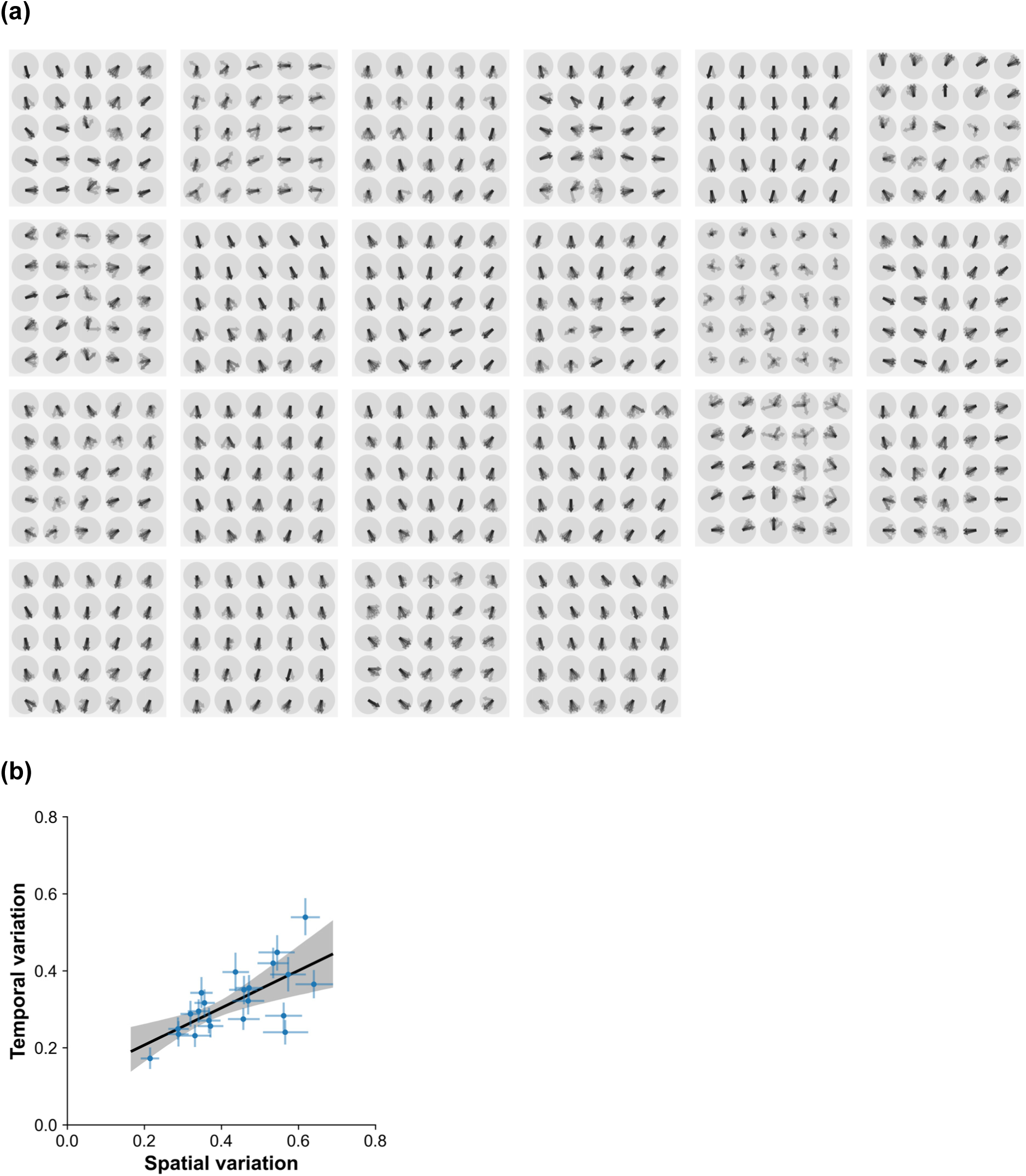
Spatio-temporal interactions. (a) The biases of all 22 participants, showing variability both across space and time. The biases are shown at each of 5x5 spatial positions for each of the 8 time slices as partially transparent arrows. (b) For each participant, the mean difference between the biases (expressed as a Euclidean distance) of nearest spatial neighbors (horizontal axis) versus the mean distance between the biases of nearest temporal neighbors. The horizontal and vertical error bars on the individual points show bootstrap-derived 95% confidence intervals. The black line is the linear regression through the means, the gray region the 95% confidence interval of the regression, derived by bootstrap. The The

I calculated the Euclidean differences between bias vectors and their nearest neighbors in space (but at the same moment in time), and the differences between biases and their nearest neighbors in time (but at the same position). These differences vary considerably across participants: the mean spatial difference varies from 0.21 to 0.64, and the mean temporal difference from 0.17 to 0.54 (see Fig. 8b). Was there a specific correlation between these spatial and temporal variations across participants? Being interested in a specifically spatio-temporal correlation rather than the general degree of variability, I calculated the partial correlation across participants between the spatial and temporal variabilities, partialling out the general variability defined as the mean difference between all pairs of the 200 biases (25 positions x 8 times) for each participant. The partial correlation coefficient between spatial and temporal was found to be +0.781, with a 95% confidence interval of [10.387, 0.936] (bootstrap over participants with 10^5^ samples). Thus, participants who had larger bias variations across spatial nearest neighbors also had larger variability across temporal nearest neighbors—irrespective of any correlation to general variability.

#### Do biases vary across space and time at the same points in spacetime?

Is variability across space associated with variability in time in a more granular way than just the participant-by-participant correlation found above? I checked whether locations in the space-time grid that had higher variabilities in bias as compared to their neighbors in space also had higher variabilities in time and vice versa, *within* each participant’s data. Concretely the spatial variability was defined as

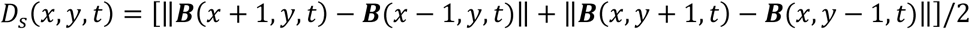

and the temporal variability as

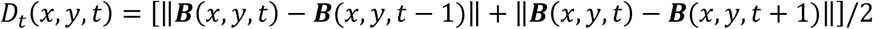

where ***B***(*x*, *y*, *t*) is the bias vector at grid position (*x*, *y*) and time *t*, and ‖… ‖ is the Euclidean distance.^6^ The Pearson correlations of *D*_s_ and *D*_t_ within each participant’s data ran from −0.241 to +0.434, with 14 out of 22 participants having positive correlations. The variance-weighted grand average of the Z-transformed correlations was 0.135, with a 95% confidence interval of [0.040, 0.223] (bootstrap over participants with 10^5^ resamples). A randomization test in which each participant’s bias field was scrambled in space and time revealed that the grand average correlation was significantly positive (*p* < 0.001 , 2-sided, 10^4^ randomizations per participant). Thus, the correlation between spatial and temporal variation holds at a quite specific level: points in space and time where the bias varies more over space tend to also be where bias varies more over time.

It turns out, however, that both spatial and temporal variabilities have small but significant negative correlations with a third variable, bias strength (bootstrap confidence intervals [−0.257, −0.007] and [−0.386, −0.143], respectively): weaker biases vary more both in space and in time. (I thank an anonymous reviewer for suggesting this analysis.) To test whether spatial and temporal variabilities are directly related, rather than through this third variable, I calculated their partial correlations, factoring out their dependence on bias strength. The result was a smaller overall correlation, 0.088, with a 95% confidence interval of [0.000, 0.167], placing it at the edge of statistical signficance. A randomization test in which biases were scrambled in space and time, however, indicated that this correlation was significantly positive (*p* = 0.014, 2-sided).

## Discussion

By measuring idiosyncratic biases on a spatial grid over the visual field and at multiple times, this study has shown that, in most cases, biases vary both over space and over time. Biases not only vary between observers as we have shown in the past (Wexler, 2018; Wexler et al., 2022, 2015), but the way that the biases vary across spatial positions in the visual field is also different from one observer to the next. Moreover, and although there are large idiosyncratic differences in bias spatial patterns, when averaged across observers the mean emergent bias pattern seems to reflect the surface-seen-from-above prior (Mamassian and Landy, 1998), not just in its value, but in its variations across space.

Importantly, this study has also shown that biases at different points in space and time are not independent of one another, but follow specific regularities. In particular, biases are spatially autocorrelated: the closer two points are in space, the more similar the biases at those two points on the average. A similar autocorrelation holds over time: the closer two bias samples are in time, the more similar the biases to each other. Furthermore, spatial and temporal variations are not independent: larger spatial variations are accompanied by greater variation over time, both participant-by-participant, and within participants point-by-point in spacetime.

These results highlight the fact that perceptual biases, even though they vary less over time than they do from observer to observer and across spatial positions within observers, are not static parameters. Instead, all evidence points to the fact that biases are dynamic, psychophysical measurements of internal brain states. Previous evidence has shown that these states, when these states are measured multiple times, they do not exhibit independent random variations about a central value. Instead, they evolve in a lawful way, acting as memories that accumulate changes over time (Wexler, 2018; Wexler et al., 2015). The new evidence presented here shows that these states show variation with non-trivial structure over space as well, and correlations between spatial and temporal variation.

What can be the neural mechanisms of the spatial and temporal fluctuations in the biases, and of the regularities in these fluctuations? Another context where visual effects are spatially autocorrelated is the spreading of visual adaptation to unadapted parts of the visual field (Altan and Boyaci, 2020; Gurbuz and Boyaci, 2023). The degree of adaptation appears to vary continuously as a function of location in the visual field: it is spatially autocorrelated. A plausible mechanism for such spreading is a visual hierarchy of brain areas with different receptive field sizes (Altan and Boyaci, 2020; Gurbuz and Boyaci, 2023). If such a hierarchy is ‘well-ordered’, meaning that a neuron with a large receptive field received input from and provides feedback to neurons with smaller receptive fields in a spatially contiguous region, such a mechanism is likely to guarantee spatial autocorrelation. The spatial autocorrelation of biases found here may indicate that a hierarchical network with different receptive field sizes is involved in the maintenance and temporal evolution of visual biases.

What sorts of neural mechanisms can lead to correlations between spatial and temporal variations in biases? Historically, Westheimer, Köhler and other Gestalt psychologists thought of the cortex as a physical medium governed by field equations, and that the neural dynamics that followed these field equations explained various perceptual phenomena such as apparent motion. A modern version of these theories are the current neurophysiological and psychophysical studies that measure spatiotemporal waves in cortex and their perceptual effects (Chavane et al., 2000; Dugué and Chavane, 2025; Muller et al., 2018). Many common field equations, such as the wave and diffusion equations, are precisley statements of the covariation between spatial and temporal variabilities at different orders (Morse and Feshbach, 1953). The finding of the correlation between spatial and temporal variations in the biases in the present study may indicate spatiotemporal field-like behavior in the brain circuits that are responsible for visual biases.

Another, somewhat peripheral and unexpected finding of the present study deserves comment: the emergent ‘grand-mean’ spatial pattern of biases when averaging over observers and over time (Figs. 6a, b). The view-from-above bias has long been documented (Mamassian and Landy, 1998), and in the context of structure-from-motion biases corresponds to the fact that a majority of the population (but far from everyone) has a preferred tilt close to the ground plane (Wexler et al., 2015). What is new is that the population mean spatial map of biases seems to reflect the variations in observer-relative surface tilt of the ground plane. This extends a result that showed that the view-from-above bias in the perception of the Necker cube is modulated by position (Dobbins and Grossmann, 2010). A remarkable aspect of the present result is that individual spatial maps do not resemble the spatial variations of the tilt of the ground plane; only when averaging over 19 observers and 8 time slices do we obtain such a map. This indicates that the large inter-observer and temporal variations of bias may actually fluctutae about a central attractor that incorporates the three-dimensional geometry of the ground plane.

There are multiple reasons to study idiosyncratic perceptual biases and their variations over observers, space, and time. Most obvious is the fact that they strongly influence perception. Here we have studied their influence on the perception of 3D orientation from ambiguous structure-from-motion stimuli, but it has been shown that similar biases operate on many different families of visual stimuli (Afraz et al., 2010; for example see Carter and Cavanagh, 2007; Wexler, 2018; Wexler et al., 2022, 2015) and even auditory stimuli (Deutsch, 1986). These differences can, among other things, cause different observers to perceive the same stimulus in opposite ways. Another reason, perhaps just as important, is that these states exist and seem to be an important but hithertofore neglected part of percesptual systems’ mechanisms. A more complete understanding of how they evolve and interact with ongoing perceptual processes is required for an fully adequate picture of perceptual mechanisms.

## Footnotes

1 The same machine-learning analysis on *temporal* patterns was less successful. Training the classifier to predict the participant from the temporal evolution of the bias at all times and all-but-one spatial locations, and testing at the untrained position, yields a generalization performance of only 41.1%: much better than the chance level of 1/18 = 5.6%, but considerably worse that the spatial analysis.

2 A field *φ* is continuous at point *s* if *φ*(*r*) approaches φ(*s*) as *r* approaches *s*.

3 *Nearest neighbors* in space will be defined as points that lie to the left, right, above and below a given point (but not diagonals), at the same time moment in time; thus, a point may have 4, 3, or 2 nearest neighbors in space. Analogously, nearest neighbors in time will be points at the same spatial position, but just before or just after in time; thus, a point may have 2 or 1 nearest neighbors in time.

4 As the variance decreases, autocorrelations should decrease and become dominated by measurement noise. Weighting by variance ensures a smaller contribution from time slices that have little variance and therefore noisy autocorrelation centered at zero.

5 The reader might wonder how an autocorrelation of +0.05 could be significantly positive. This is because, counterintuitively, the expectation value of the autocorrelation of a series with independent random terms is not zero but negative. For example, the expectation value of a series with 8 terms independently chosen from a semicircular distribution (like the components of a random bias inside the unit disk) is about −0.14. It turns out that the autocorrelation estimate of +0.085is about two standard deviations above its negative mean in the null distribution.

6 Other choices could have been made here, and yield similar results. An important point is to avoid common terms like ***B***(*x*, *y*, *t*) in both *D*_s_ and *D*_t_, which would lead to spurious correlations.

## Notes

### Competing Interest Statement

The authors have declared no competing interest.

