## Supplemental figure 1 for "Dynamics of individual visual biases in space and time"

### Supplementary materials

**Figure S1.** Raw data from all 52,800 trials, shown together with biases. Each page-wide row surrounded by a gray border represents one participant (total of 22 participants). Within each row, each square represents one of 8 time slices, approximately 15 minutes. Within each square, each circle represents one of 5x5 spatial positions, separated by 5.75 degrees of visual angle. Within each position, each of the 12 trials is shown as two opposite-pointing arrows: the green arrow shows the reported tilt, while the pink arrow shows the tilt that wasn't reported (see Figs. 1e,f). The black arrow shows the bias for that spatio-temporal location; its direction shows the preferred tilt, and its length represents bias strength (see Fig. 1g,h).

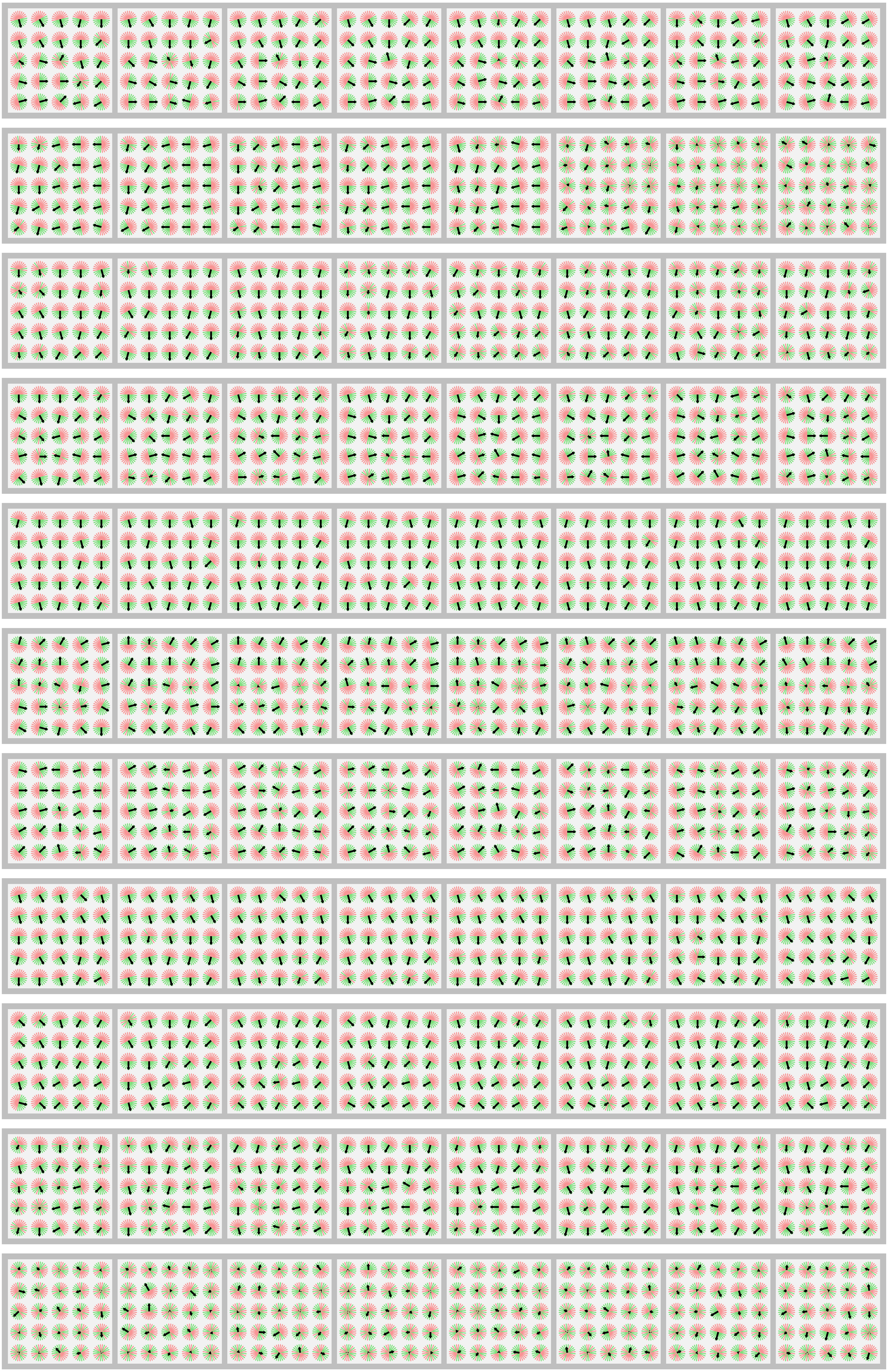

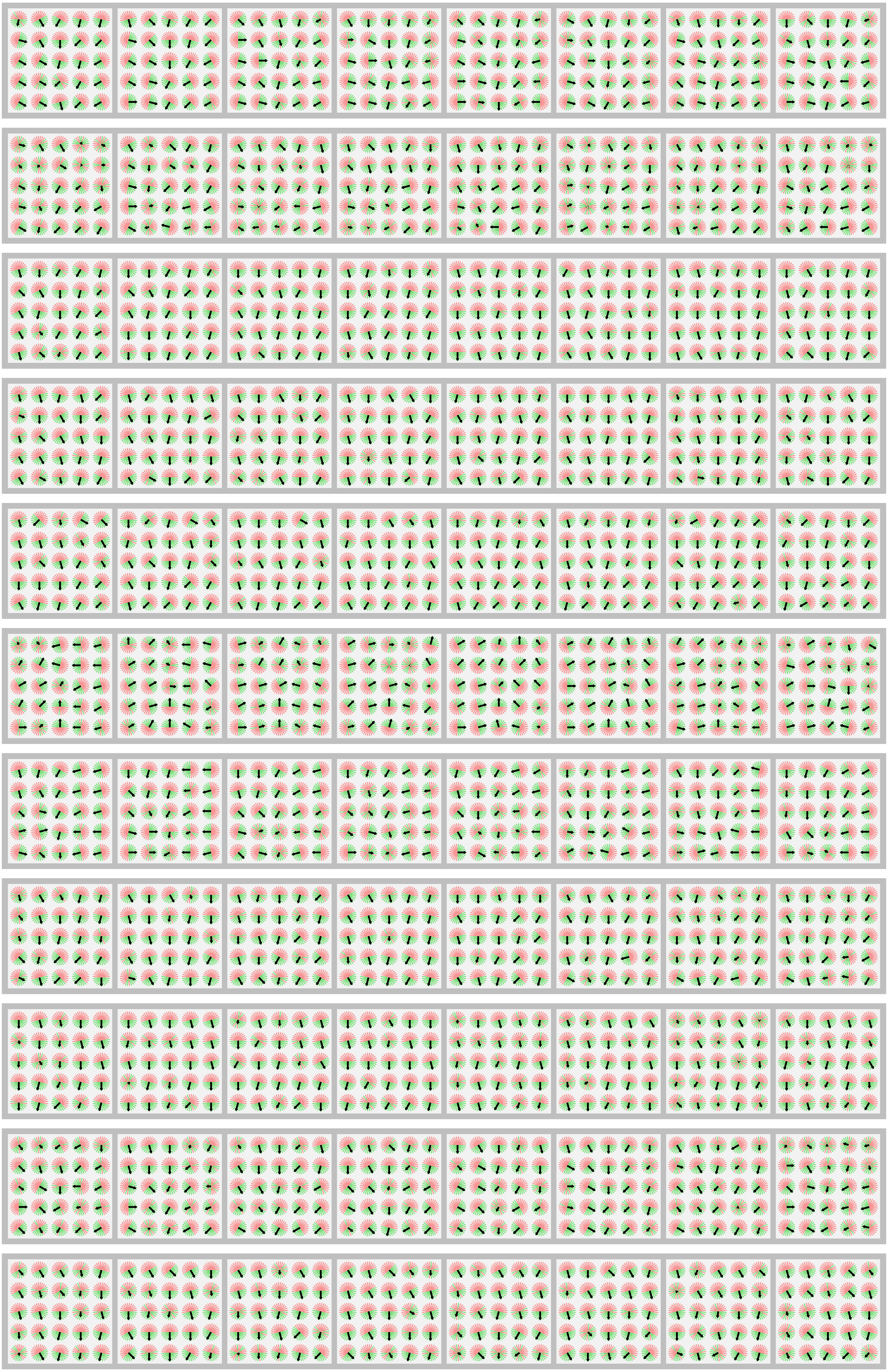
